# Global profiling of the crotonylome in Small Cell Lung Cancer

**DOI:** 10.1101/2020.06.29.175877

**Authors:** Zongpei Guo, Meng Gu, Jiaqiang Huang, Ping-kun Zhou, Teng Ma

## Abstract

Small cell lung cancer is a deadly neuroendocrine lung cancer subtype, which harbors the driver mutations of *RB1* and *TP53* as well as *EP300/CREBBP*. Since CREBBP/p300 are two prominent crotonyl-CoA transferases responsible for the global crotonylation in cells, here we investigated the crotonylome in SCLC tissues compared with normal lung tissues through TMT labeling coupled with LC-MS/MS and found unique patterns of protein crotonylation in SCLC related pathways.

## Materials and Methods

### Cell culture

H526 cells purchased from ATCC were maintained in ATCC-formulated RPMI-1640 Medium supplemented with 10% FBS. Cells were cultured at 37°C with 5% CO_2_.

### Patient samples

Paired SCLC tumor and normal tissues were purchased from Shanghai SuperChip Biotech Co. Ltd.

### Antibodies

Commercial antibodies were used as follows: anti-STMN1 (Abcam, ab52630), anti-crotonyllysine antibody (PTM, 501), anti-β-actin (Abcam, ab8227).

### Protein extraction and trypsin digestion

The tissue samples are taken from −80 °C and ground thoroughly to powder with liquid nitrogen. Then 4X volumes of lysis buffer was added to the tissue powder (8M urea, 1% protease inhibitor, 3μM TSA and 50mM NAM). Samples were sonicated. The remaining debris was removed by centrifuge at 12,000g at 4□ for 10min. The supernatant was collected and the protein concentration was determined with a BCA kit according to the manufacturer’s instructions. For trypsin digestion, the protein supernatant was treated with 5mM dithiothreitol (DTT) for 30min at 56°C and alkylated with 11mM iodoacetamide for 15min at room temperature in darkness. The protein samples were then diluted with 100mM triethylammonium bicarbonate (TEAB) to urea concentration of less than 2M. Finally, trypsin was added at 1:50 mass ration of trypsin/protein and proteins were digested overnight. A second 4h digestion was performed with trypsin/protein at 1:100 mass ration.

### TMT labeling

After trypsin digestion, peptide was desalted by Strata X C18 SPE column (Phenomenex) and vacuum-dried. Peptide was reconstituted in 0.5 M TEAB and processed according to the manufacturer’s protocol for TMT kit. Briefly, one unit of TMT reagent were thawed and reconstituted in acetonitrile. The peptide mixtures were then incubated for 2 h at room temperature and pooled, desalted and dried by vacuum centrifugation.

### Crotonylation modified peptide enrichment

To enrich modified peptides, tryptic peptides dissolved in NETN buffer (100 mM NaCl, 1 mM EDTA, 50 mM Tris-HCl, 0.5% NP-40, pH 8.0) were incubated with pre-washed antibody beads (Lot number 001, PTM Bio) at 4°C overnight with gentle shaking. Then the beads were washed four times with NETN buffer and twice with H_2_O. The bound peptides were eluted from the beads with 0.1% trifluoroacetic acid. Finally, the eluted fractions were combined and vacuum-dried. For LC-MS/MS analysis, the resulting peptides were desalted with C18 ZipTips (Millipore) according to the manufacturer’s instructions.

### LC-MS/MS analysis

The tryptic peptides were dissolved in 0.1% formic acid (solvent A), directly loaded onto a home-made reversed-phase analytical column (15-cm length, 75 μm i.d.). The gradient was comprised of an increase from 6% to 23% solvent B (0.1% formic acid in 98% acetonitrile) over 26 min, 23% to 35% in 8 min and climbing to 80% in 3 min then holding at 80% for the last 3 min, all at a constant flow rate of 400 nL/min on an EASY-nLC 1000 UPLC system.

The peptides were subjected to NSI source followed by tandem mass spectrometry (MS/MS) in Q Exactive™ Plus (Thermo) coupled online to the UPLC. The electrospray voltage applied was 2.0 kV. The m/z scan range was 350 to 1800 for full scan, and intact peptides were detected in the Orbitrap at a resolution of 70,000. Peptides were then selected for MS/MS using NCE setting as 28 and the fragments were detected in the Orbitrap at a resolution of 17,500. A data-dependent procedure that alternated between one MS scan followed by 20 MS/MS scans with 15.0s dynamic exclusion. Automatic gain control (AGC) was set at 5E4. Fixed first mass was set as 100 m/z.

### Database search

The resulting MS/MS data were processed using Maxquant search engine (v.1.5.2.8). Tandem mass spectra were searched against human uniprot database concatenated with reverse decoy database. Trypsin/P was specified as cleavage enzyme allowing up to 4 missing cleavages. The mass tolerance for precursor ions was set as 20 ppm in First search and 5 ppm in Main search, and the mass tolerance for fragment ions was set as 0.02 Da. Carbamidomethyl on Cys was specified as fixed modification and Acetylation modification and oxidation on Met were specified as variable modifications. FDR was adjusted to < 1% and minimum score for modified peptides was set > 40.

### Bioinformatics analysis

Gene Ontology (GO) annotation proteome was derived from the UniProt-GOA database (http://www.ebi.ac.uk/GOA/). Identified proteins domain functional description were annotated by InterProScan (a sequence analysis application) based on protein sequence alignment method, and the InterPro (http://www.ebi.ac.uk/interpro/) domain database was used. Kyoto Encyclopedia of Genes and Genomes (KEGG) database was used to annotate protein pathway. We used wolfpsort, a subcellular localization predication soft to predict subcellular localization. Soft MoMo (motif-x algorithm) was used to analysis the model of sequences constituted with amino acids in specific positions of modify-21-mers (10 amino acids upstream and downstream of the site). Enrichment of GO or KEGG analysis was done by a two-tailed Fisher’s exact test to test the enrichment of the differentially modified proteins. STRING database version 10.1 was used for protein-protein interactions and Cytoscape software was used for visualization.

### Immunoprecipitation and western blotting

For immunoprecipitation, cells were lysed with NETN-300 buffer (20mM Tris-HCl Ph8.0, 300mM NaCl, 1mM EDTA, 0.5% Nonidet P-40) containing protease inhibitor cocktail (Roche), 3μM TSA and 50mM NAM on ice for 10min, then supplemented double volume of NETN-100 buffer (20mM Tris-HCl pH8.0, 100mM NaCl, 1mM EDTA, 0.5% Nonidet P-40). The lysates were centrifuged at 12000g for 10min at 4□. The supernatants were collected and incubated with 1.5μg anti-STMN1 antibody overnight at 4□, then protein A/G agarose (Santa Cruz) was added and incubated for 3h at 4□. Then the agarose was collected by centrifuging at 1000g for 3min and washed 3 times by NETN-100 buffer, and the immunoprecipitated proteins detected by western blot.

### Main text

Small Cell Lung Cancer accounts for ~15% of lung cancer incidence with highest mortality. Concurrent driver mutations of *RB1* and *TP53* have been clearly demonstrated^1,2^. Mutual exclusive mutations of *CREBBP*/*EP300* have also been found. The CREBBP and p300 proteins as well as PCAF and MOF catalyze the crotonylation on histones and non-histone proteins^3^. Crotonylation has been demonstrated to be involved transcription and DNA repair^3,4^. Though several proteomic studies have been focused on the regulation of crotonylation in cells^4–6^, the relationship between protein crotonylation and cancer is unknown. Here we collected resected small cell lung cancer tissues from 3 different patients along with the paired normal lung tissue as control. The TMT labeling and LC-MS/MS analysis was shown in flowchart (Figure 1A). We finally identified 1712 sites of 662 proteins containing quantitative information. The data was normalized with a protein quantification proteomic to exclude the effect of protein expression on the modification signal.

**Figure 1.**
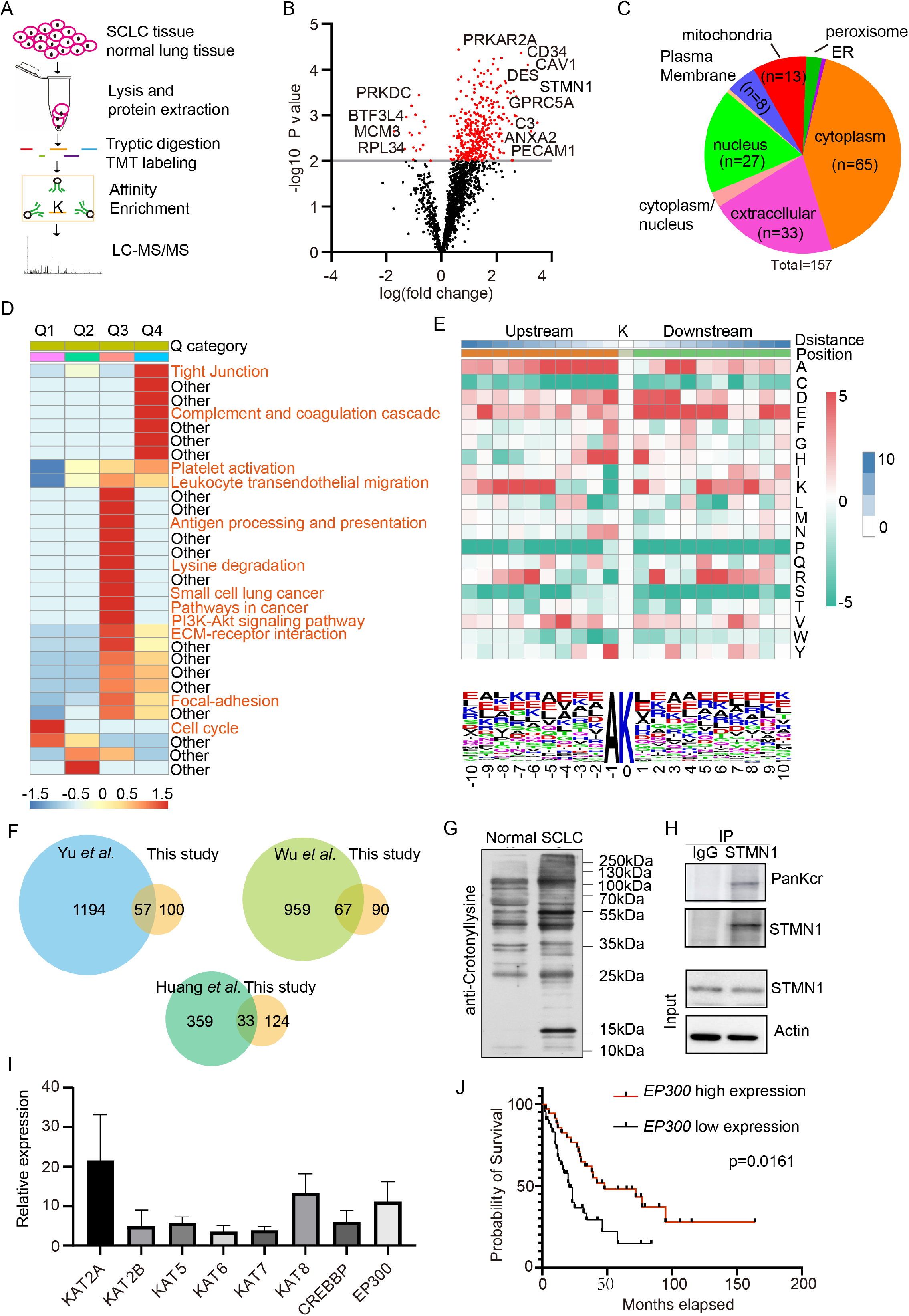
Identification of crotonylome in SCLC. (A) Flow chart of the proteomics procedure. (B) The volcano distribution of differentially modified proteins. (C) The subcellular distribution of differentially modified proteins. (D) The 4 categories cluster of KEGG pathway analysis of differentially modified proteins. (E) Motif analysis of all identified crotonylated sites. (F) Venn Diagram of the overlapping crotonylated proteins between previous studies and this study. (G) Overall crotonylation detection in SCLC tissue and normal control. (H) Validation of STMN1 as crotonylated protein in H562 cells. (I) Relative expression profile of known crotonyl-CoA transferases. (J) Kaplan-Meier analysis of survival probability of SCLC patients with either high or low *EP300* expression.

The differentially modified sites between SCLC tissue and normal control were screened following the criteria: 2 times as the change threshold and t-test *p*-value < 0.05. The Number of modification sites per protein and peptides length are shown in Figure S1A and S1B. The distribution of mass errors was near zero indicating accuracy of the MS data (Figure S1C). Based on the above data and criteria, we found 368 upregulated modified sites of 145 proteins in the SCLC tissues compared with Normal control, and 14 downregulated sites of 13 proteins. (Figure 1B and Figure S1D). As shown in volcano plot of Figure 1B, several important regulators of tumor metastasis or tumor microenvironment such as PECAM1 (Platelet and Endothelial Cell Adhesion Molecule 1; CD31)^7^, CAV1 (caveolin-1)^8–10^, Complement C3^11,12^ and STMN1 ^13^ are modified with crotonylation in SCLC tissues. The protein subcellular analysis showed that 65 proteins were in cytoplasm, 33 proteins in extracellular and 27 proteins in nucleus (Figure 1C), suggesting a broad spectrum of functions of crotonylation modified proteins in SCLC.

Next, Gene Ontology classification and enrichment analysis showed that crotonylation modified proteins in SCLC are involved processes of wound healing which is the hallmark of cancer (Figure S2) ^14,15^. To gain an insight of crotonylome function in SCLC, we divided the differentially modified sites into 4 categories based on the change ration of SCLC/Normal, namely Q1 (<0.05), Q2 (0.500-0.667), Q3 (1.5~2) and Q4 (>2). Then each category was clustered according to KEGG analysis as it showed in Figure 1D. In Q4, tight junction^16^, complement and coagulation and Platelet activation^17^ were most related pathways to SCLC or cancer. In Q3 cluster, both SCLC signature and Lysine degradation pathway which produces crotonyl-CoA were found. Immune regulation pathways and cancer related pathways such as PI3K-Akt were also found (Figure 1D). Protein domain enrichment analysis indicated that the EF-hand domains are closely related with SCLC and inflammation (Figure S3)^18^. The PPI network indicated the SCLC crotonylome falls into several complexes (Figure S3).

To investigate the amino acid frequency of occurrence upstream and downstream from the crotonylation modification site, the flanking sequences were analyzed. The results showed that alanine (A) and glutamate (E) residues were overrepresented at the −1 and +1 positions surrounding the crotonylated lysine respectively (Figure 1E). Further characterization of the motifs surrounding lysine crotonylation sites showed a series of different patterns of crotonylation motifs and a representative motif logo was shown (Figure 1E).

To investigate whether the crotonylation modified proteins overlap with previous cell line-based proteomics studies, the differentially modified proteins in SCLC vs Normal from our study were compared with Yu et al. study which is about CDYL regulated crotonylome, Wu et al. study which is about non-histone crotonylome and Huang et al. study which is about p300 regulated crotonylome. 57, 67 and 33 proteins were found to overlap with their findings respectively (Figure 1F). There are some common targeted proteins such as STMN1 and histones in both p300 regulated crotonylome and SCLC/normal crotonylome. However, the majority modified proteins are not over-lapping, highlighting the unique crotonylome in SCLC tissue from previous cellular proteomics. Furthermore, the pan-crotonylation signal was increased in SCLC tissue vs normal control (Figure 1G). STMN1 was modified by crotonylation (Figure S4) and validated in H526 cells (Figure 1H).

Since CREBBP, p300, PCAF(KAT2B) and MOF(KAT8) are the major crotonyl-CoA transferases in cells, we investigated their relative expression profiles as well as other known acetyl-CoA transferases in SCLC tissues. RNA-Seq data of 81 SCLC patients from George et al. study were extracted and analyzed^2^. *KAT2A (GCN5)*, *KAT8 (MOF)* and *EP300* showed relative higher expressions in SCLC. Next we checked whether any of these crotonyl-CoA transferases could associate with patients prognosis. The results demonstrated that only p300 had the favorable overall survival in the Kaplan-Meier survival analysis with *P* value =0.0161 while *KAT2A*, *KAT8* and *CDYL* (the crotonylation hydratase) had no prognostic significance (Figure S5), suggesting that p300’s function as crotonyl-CoA transferase could play tumor suppressor functions in SCLC which is similar to the reported role of CREBBP in SCLC^16^.

In summary, our study is the first systematic analysis of crotonylation in SCLC tissues. Further investigations of the functions of protein crotonylation in diverse pathways may shed light on the understanding of malignant behaviors of SCLC including metastasis, immune suppression and chemo/radio-resistance.

## Acknowledgement

This study was supported by grants from the National Natural Science Foundation of China (31570853, 81602799, 81530085, 31870847).

