## Supplemental Material for "Global profiling of the crotonylome in Small Cell Lung Cancer"

### Supplementary material

#### Supplementary Figures and Figure legends

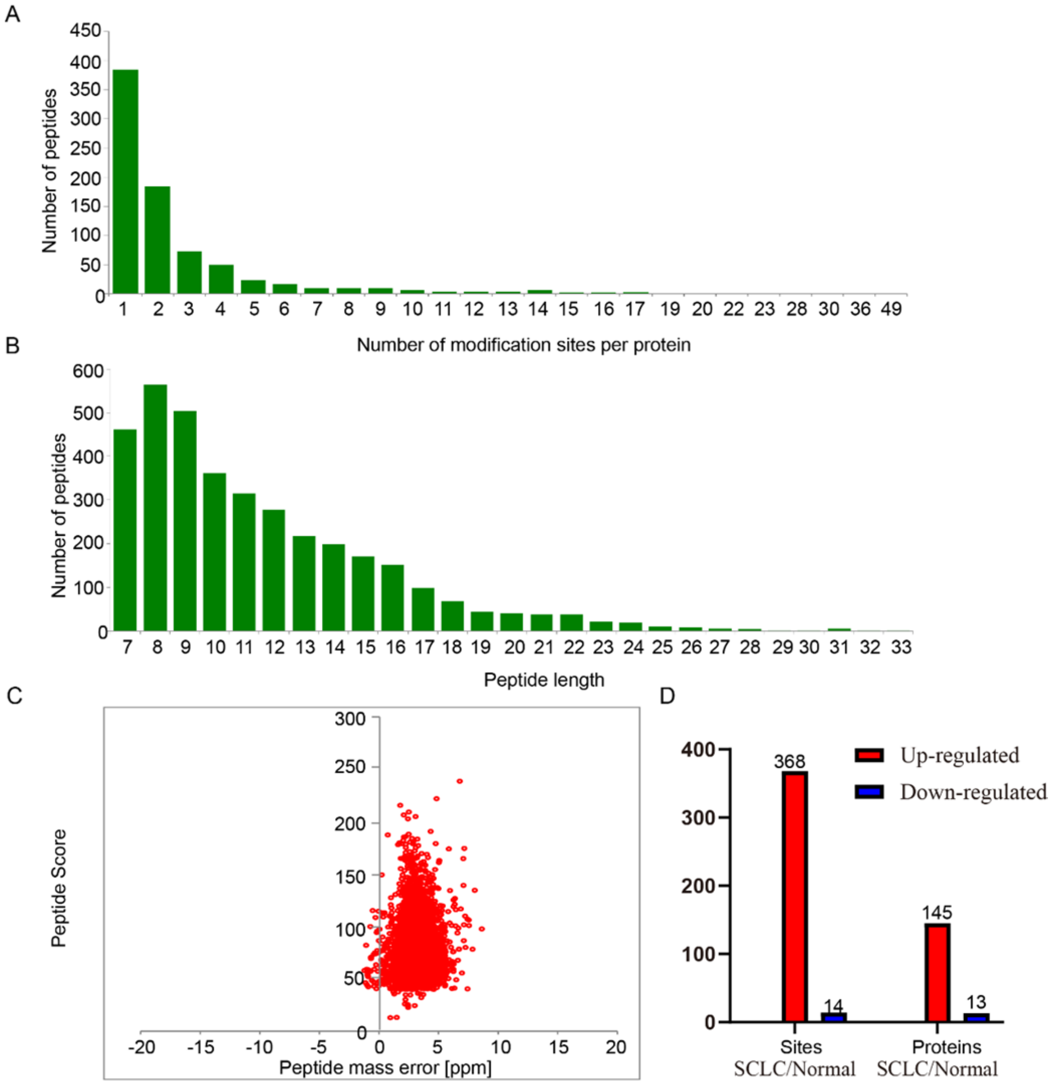


Figure S1. Identification of differential protein crotonylation in SCLC. (A) Most of the identified peptides contain less than 3 modification sites. (B) Most peptides are less than 20 amino acid in length. (C) The peptide mass error was nearly zero. (D) Summary of identified sites and proteins.

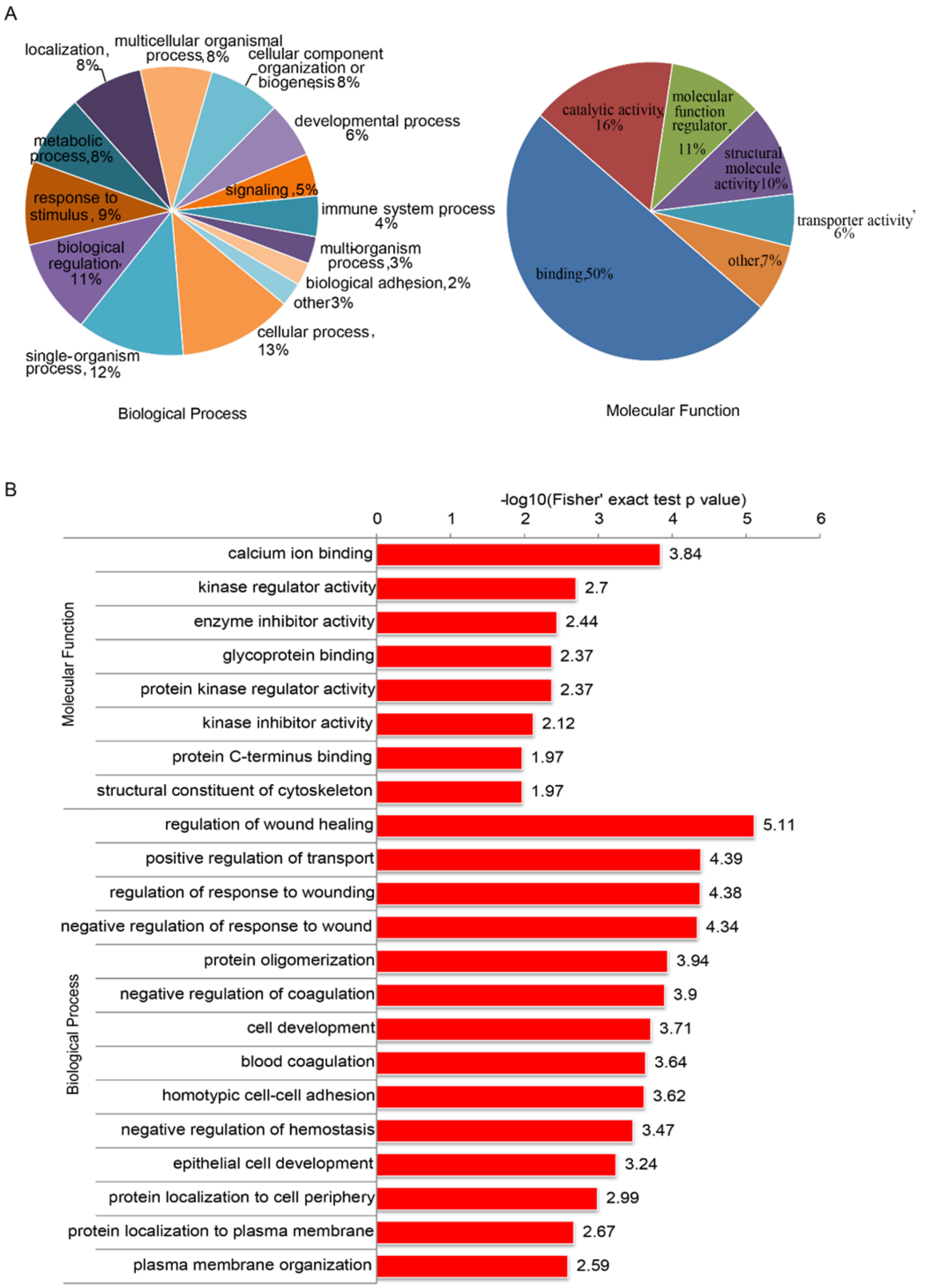


Figure S2. Go classification and enrichment analysis of the differentially modified proteins. (A) The biological process and molecular function analysis of differentially modified proteins. (B) The GO enrichment analysis after a Fisher' exact test.

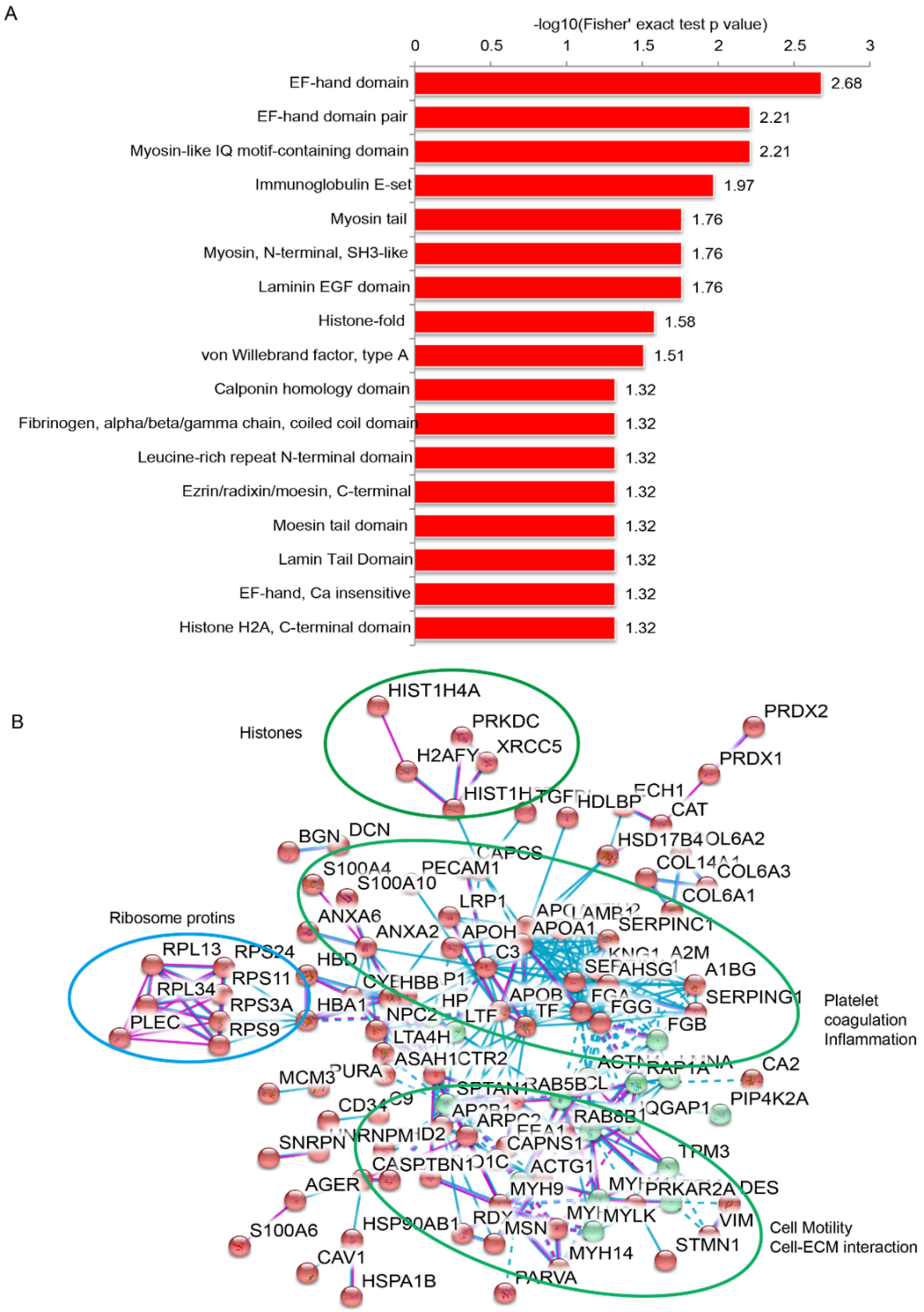


Figure S3. Go enrichment analysis of the differentially modified domains (A) and the PPI network from STRING analysis (B).

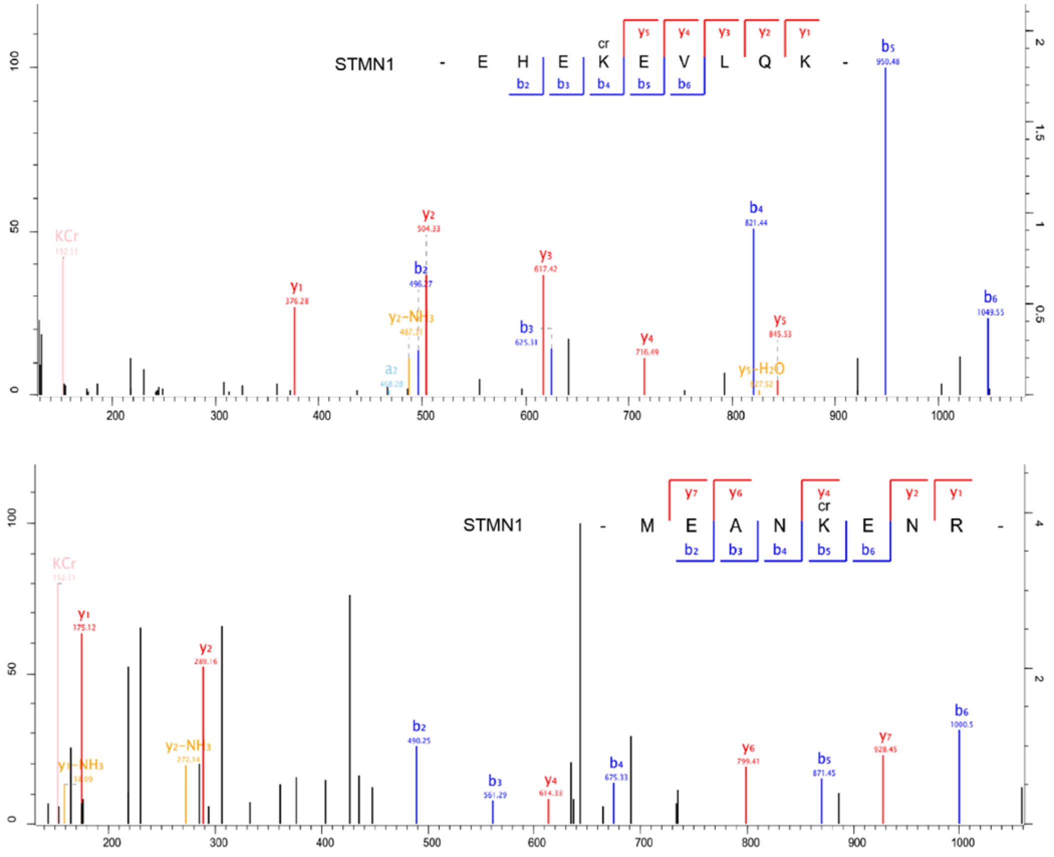


Figure S4. The mass spectrometry showed 2 differentially modified sites of STMN1.

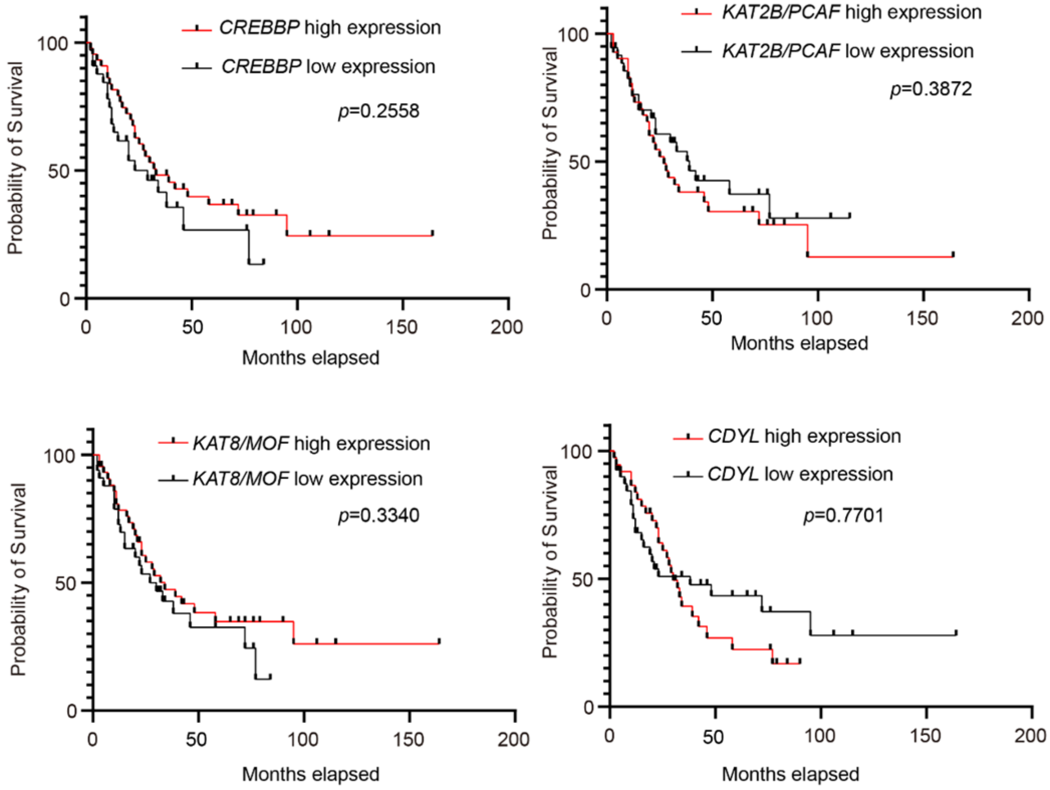


Figure S5. The Kaplan-Meier analysis of survival probability in SCLC patients with high or low expressions of *CREBBP*, *KAT2B/PCAF*, *KAT8/MOF* and *CDYL*.
